# Different speciation types meet in a Mediterranean genus: the biogeographic history of *Cymbalaria* (Plantaginaceae)

**DOI:** 10.1101/050369

**Authors:** Pau Carnicero, Llorenç Sáez, Núria Garcia-Jacas, Mercè Galbany-Casals

**Affiliations:** Departament de Biologia Animal, Biologia Vegetal i Ecologia, Facultat de Biociències, Universitat Autònoma de Barcelona, 08193 Bellaterra, Spain.; Societat d’Història Natural de les Balears (SHNB), C/ Margarida Xirgu 16, E-07011 Palma de Mallorca, Balearic Islands, Spain.; Institut Botànic de Barcelona (IBB-CSIC-ICUB), Pg. del Migdia s/n, ES-08038 Barcelona, Spain

**Keywords:** Ancestral-area estimation, cpDNA, founder-event speciation, long-distance dispersal, molecular dating, nrDNA

## Abstract

*Cymbalaria* is a Mediterranean genus including ten species and six subspecies growing on rocky habitats. A very fragmented distribution, the different ecologic preferences of partially sympatric species and the presence of different ploidy levels, suggest the role of allopatric, sympatric ecological and polyploid speciation in its evolution. The aims of this study are to verify the monophyly and to reconstruct the phylogenetic relationships of *Cymbalaria*, to infer its biogeographic history by estimating the lineage divergence dates and the ancestral areas of distribution, and to discuss the role of different types of speciation. To address these issues, we constructed a complete phylogeny of the genus with ITS, 3’ETS, *ndh*F and *rpl32-trn*L sequences. A time-calibrated phylogeny and an ancestral-area estimation were obtained from the nrDNA data. The evidence supported the genus *Cymbalaria* as monophyletic. It originated ca. 5 Ma and three lineages segregated rapidly, one with the single extant taxa *Cymbalaria microcalyx* subsp. *microcalyx* and the other two corresponding to western and central-eastern species, respectively. The main diversification events occurred after the onset of the Mediterranean climate and during Pleistocene oscillations. Founder-event and sympatric speciation were supported by the biogeographic analyses, and chromosome data combined with our phylogeny supported at least two polyploidization events. We observed that the consequences of physical barriers were different amongst the different species.

**Supplementary Material:** Electronic Supplement (Figure S1, Table S1) are available in the Supplementary Data section of the online version of this article (http://www.ingentaconnect.com/content/iapt/tax).

## INTRODUCTION

The Mediterranean Basin contains ca. 25,000 species and almost 10% of the world’s vascular flora, of which 63% are endemics (Greuter, 1991). Three primary types of speciation might have originated this high diversity (Thompson, 2005). First, allopatric speciation is favoured in a fragmented landscape with a history of temporary land connections and isolations among the mainland and the numerous islands. Allopatric speciation is coupled with the effects of two major climatic events: the establishment of a Mediterranean climate approximately 3.2 Ma, which marked an increase in the rates of diversification for many plant lineages (Fiz-Palacios & Valcàrcel, 2013), and the Pleistocene glaciations, which altered the distributions of species and favoured gene flow among populations in some cases, whereas in other cases populations became isolated in climatic refuges (Vargas, 2003; Médail & Diadema, 2009). Second, sympatric ecological speciation has also been documented (Santos-Gally & al., 2011) and is favoured by the wide heterogeneity of habitats and the altitudinal gradients in relatively small areas. Third, polyploid speciation is proposed for many Mediterranean plant groups (Thompson, 2005). Because of the increase in genetic variation, polyploids are successful in the colonization of new niches to expand their ranges (Ramsey, 2011).

*Cymbalaria* Hill. (Plantaginaceae) is a genus of perennial herbs with ten species and six subspecies (Sutton, 1988; Bigazzi & Raffaelli, 2000), distributed throughout the Mediterranean Basin (Fig. 1). *Cymbalaria muralis* G. Gaertn., B. Mey. & Scherb., although native to the central Mediterranean Basin, is naturalised almost worldwide in temperate areas (Sutton, 1988) and is therefore the most widespread species. The last complete systematic revision of the genus was carried out by Sutton (1988), in which he highlighted some taxonomic conflicts, mainly regarding eastern Mediterranean taxa. *Cymbalaria* has been included in molecular studies on the tribe Antirrhineae (Ghebrehiwet & al., 2000; Vargas & al., 2004, 2013; Guzmán & al., 2015), but molecular analyses with a comprehensive sampling of the genus have never been performed. All *Cymbalaria* species grow in rocky habitats in a wide range of ecological conditions, from coastal cliffs to rock crevasses in the subalpine stage. The rocky habitats and most of the areas currently occupied by *Cymbalaria* species are considered to have remained climatically stable during Pleistocene glaciations (Thompson, 2005; Médail & Diadema, 2009), suggesting an important role of climatic refugia in the evolutionary history of *Cymbalaria*. Geographic isolation might have played a different role in speciation, since some species are narrow endemics geographically isolated from other taxa, while other species show very fragmented, disjunct, but broad distribution areas (Fig. 1). The last pattern could be caused by recent range expansion or by active gene flow among disjunct populations. Some species occur in sympatric areas but with well differentiated ecological preferences, likely suggesting the action of sympatric ecological speciation (Fig. 1, Table 1). Ploidy levels vary across species and are often geographically grouped, ranging from diploids (2*n* = 14) to octoploids (2*n* = 56, Fig. 1), supporting an important impact of polyploidy in driving speciation. Diploids mainly occur in the Apennine and Balkan peninsulas, with one species in the eastern Mediterranean; tetraploids (2*n* = 28) occur in Sicily, the Balkan Peninsula and the eastern Mediterranean basin, and a group of hexa-to octoploids (2*n* = 42, 56) occur in Corsica, Sardinia and the Balearic Islands. The aforementioned features make *Cymbalaria* an exemplary case for the study of plant speciation in the Mediterranean Basin, suggesting that several processes and types of speciation originated the current diversity and distribution.

**Table 1.** Chromosome number and ecology of the 16 sampled taxa.

| **Taxon** | **Chromosome number** | **Ecology** |
| --- | --- | --- |
| *C. aequitriloba* (Viv.) A. Chev. | $2n = 42, 56^1$ | Coastal and inland shadowed cliffs, moist rocks on stream banks |
| *C. fragilis* (Rodrig.) A. Chev. | $2n = 56^2$ | Coastal and inland shadowed cliffs |
| *C. glutinosa* Bigazzi & Raffaelli subsp. *glutinosa* | $2n = 14^3$ | Coastal and inland shadowed cliffs, walls |
| subsp. *brevicalcarata* Bigazzi & Raffaelli | $2n = 14^3$ | Coastal and inland shadowed cliffs, walls |
| *C. hepaticifolia* Wettst. | $2n = 56^1$ | High-elevation rocks, moist rocks and mountain stream banks |
| *C. longipes* (Boiss. & Heldr.) A. Chev. | $2n = 14^1$ | Coastal cliffs, rocks and walls |
| *C. microcalyx* (Boiss.) Wettst. subsp. *microcalyx* | $2n = 28^4$ | Inland shadowed cliffs, walls |
| subsp. *acutiloba* (Boiss. & Heldr.) Greuter | ? | Inland shadowed cliffs |
| subsp. *dodekanesi* Greuter | $2n = 28^5$ | Inland shadowed cliffs |
| subsp. *ebelii* (Cufod.) Cufod. | $2n = 28^6$ | Inland shadowed cliffs, walls |
| subsp. *minor* (Cufod.) Greuter | $2n = 28^1$ | Inland shadowed cliffs |
| *C. muelleri* (Moris.) A. Chev. | $2n = 42^7$ | Inland overhanging cliffs |
| *C. muralis* G. Gaertn., B. Mey. & Scherb. subsp. *muralis* | $2n = 14^1$ | Inland shadowed cliffs, walls |
| subsp. *visianii* (Jáv.) D.A. Webb | $2n = 14^3$ | Inland shadowed cliffs, walls |
| *C. pallida* Wettst. | $2n = 14^1$ | High-elevation rocks, mountain stream banks |
| *C. pubescens* (J. Presl & C. Presl) Cufod. | $2n = 28^3$ | Inland shadowed cliffs, walls |

<sup>1</sup> Sutton (1988) and references therein.<sup>2</sup> Castro & Rosselló (2006).<sup>3</sup> Bigazzi & Raffaelli (2000).<sup>4</sup> Speta (1986).<sup>5</sup> P. Carnicero, unpublished data.<sup>6</sup> Speta (1989).<sup>7</sup> Onnis & Floris (1967).

**Figure 1.**
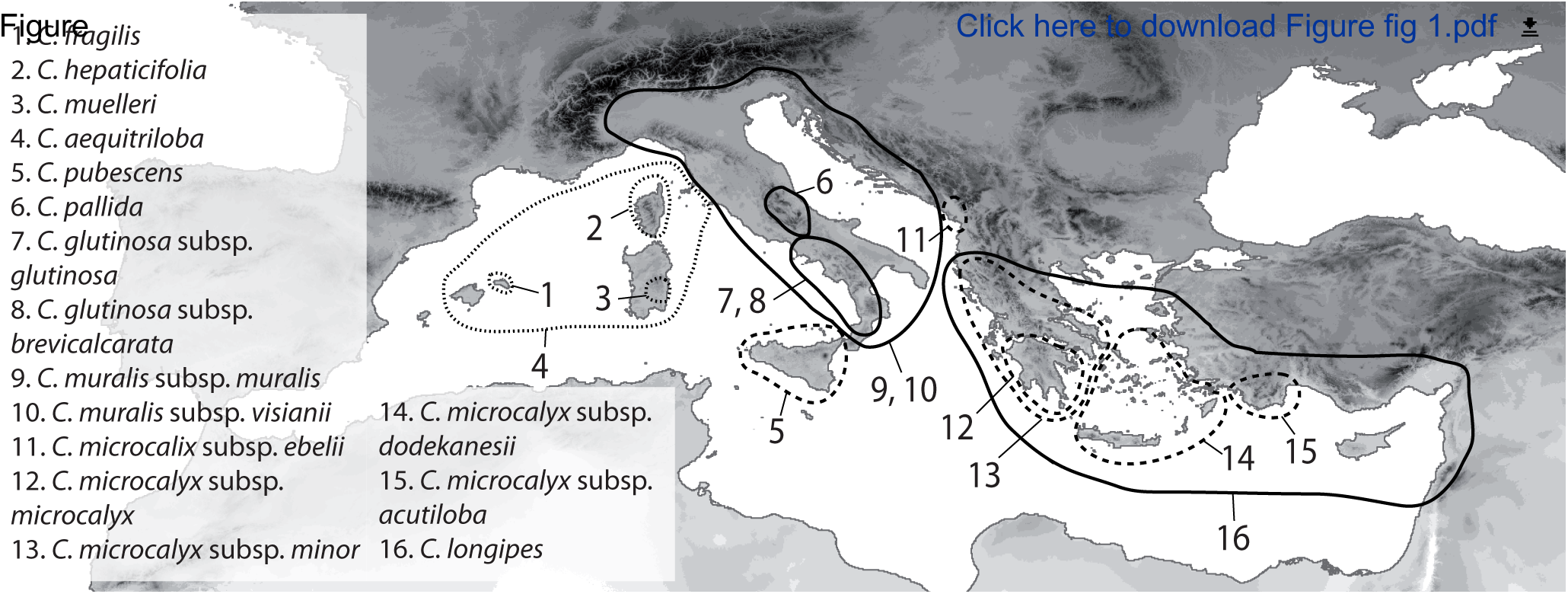
Distributions *of Cymbalaria* taxa, based on Sutton (1988), local Floras, personal field observations and herbarium vouchers. When the information on the areas of distribution of subspecies was not accurate or the overlap was considerable, the general distribution area for the species is shown. For *Cymbalaria muralis*, which is widely naturalized in temperate regions, the approximate natural distribution is shown. Different line formats indicate ploidy levels: solid line for diploids, dashed line for tetraploids and dotted line for hexa-to octoploids. Ploidy level for *C microcalyx* subsp. *acutiloba* is not known, but we assumed it to be the same as other *C microcalyx* subspecies.

Multi-locus molecular phylogenies, molecular dating, diversification analyses and ancestral area estimation models, can be used to infer the biogeographic histories of plants at different taxonomic levels (e.g. Calviño & al., 2016; Cardinal-Mc Teague & al., 2016; Janssens & al., 2016). The well-known geomorphological and climatic history of the Mediterranean Basin, together with the high interest found on studying its high plant endemism and biodiversity, make it a very suitable and attractive area for reliable reconstructions of the spatio-temporal evolution of plant lineages (e.g., Gaudeul & al., 2016; Hardion & al., 2016). Here we used cp-and nrDNA sequences to: (1) verify the monophyly of *Cymbalaria* and to clarify the phylogenetic relationships among the species, (2) estimate the divergence dates of the lineages and reconstruct the ancestral areas to infer the biogeographic history of the genus, and (3) examine the role of the different types of speciation in the evolution of *Cymbalaria*.

## MATERIALS AND METHODS

### Plant Material. –

We sampled 34 individuals of *Cymbalaria*, representing all species and subspecies recognised in the last taxonomic treatments (Sutton, 1988; Bigazzi & Raffaelli, 2000; Appendix 1). Species from 11 additional genera of the tribe Antirrhineae were also sampled to confirm the placement of *Cymbalaria* within the tribe and to assess its monophyly. *Plantago lanceolata* L. was used as the outgroup for the tribe (Olmstead & al., 2001).

### DNA extraction, amplification and sequencing. –

To extract the DNA, the CTAB method (Doyle & Doyle, 1987), as modified by Cullings (1992) and Tel-Zur & al. (1999), and the commercial kit NucleoSpin^®^ Plant were used (Macherey-Nagel GmbH & Co., KG, Düren, Germany).

We amplified the ITS region and the conserved 3’ETS region of the nuclear ribosomal DNA (nrDNA) and the *ndhF* region and the *rpl32-trnL* spacer of the chloroplastic DNA (cpDNA). We used the primers ITS1 and ITS4 (Sun & *al*., 1994) for the ITS region, Ast1 and 18SETS (Markos & Baldwin, 2001) for the 3’ETS region, 3’F (Eldenäs & *al*., 1999) and +607 (Kim & Jansen, 1995) for the *ndhF* region and rpl32F and trnL-UAG (Shaw & *al*., 2007) for the *rpl32-trnL* spacer. For some specimens, we designed and used internal specific primers for the *ndhF* region: (1) *ndhF* CymbF: 5’ TGA ATC GGA CAA TAC CAT GTT ATT 3’; (2) *ndhF* CymbR: 5’ ATT CAT ACC AAT TCG TCG AAT CCT 3’; (3) *ndhF* CymbF2: 5’ ACG AGT AAT TGA TGG AAT TAC G 3’; and (4) *ndhF* CymbR2: 5’ GAG TCT TAT CTG ATG AAT ATC 3’. The profile used for amplification of the ITS included 4 min denaturing at 95°C, followed by 30 cycles of 90 s denaturing at 94°C, 2 min annealing at 55°C and 3 min extension at 72°C, with an additional final step of 15 min at 72°C. The profile used for amplification of the *rpl32*-*trnL* spacer included 3 min denaturing at 94°C, followed by 30 cycles of 40 s denaturing at 95°C, 2 min annealing at 52°C and 2 min extension at 72°C, with an additional final step of 10 min at 72°C. We followed the PCR profiles described in Galbany-Casals & *al*. (2009) for the ETS and Galbany-Casals & *al*. (2012) for the *ndhF*. PCR products were purified with Exo-SAP-IT (USB Corp., Cleveland, Ohio, U.S.A.). Direct sequencing was conducted at the DNA Sequencing Core, CGRC/ICBR of the University of Florida, on an ABI 3730xl DNA Analyser (Applied Biosystems) using a Big Dye Terminator v.3.1 kit (Applied Biosystems, Foster City, CA, U.S.A.). See Appendix 1 and electronic supplement Table S1 for information on the vouchers and the sequences.

### Phylogenetic analyses. –

The sequences were examined and aligned by hand using Chromas Lite 2.0 (Technelysium Pty Ltd., Tewantin, Australia) and Mega 6.06 (Tamura & al., 2013). The ambiguous regions of the alignments were manually excluded. Indels were codified as binary characters using the simple indel coding method (Simmons & Ochoterena, 2000) for the cpDNA alignment. The nrDNA alignment provided enough variation so indels were not codified. Cp and nrDNA regions were analysed separately.

Maximum Parsimony (MP) analyses were conducted with PAUP*v.4.0a147 (Swofford, 2002), with 10000 replicates of heuristic searches with random taxon addition and tree bisection-reconnection (TBR) branch swapping and holding all most parsimonious trees. The indels were coded as missing data, and the uninformative characters were excluded. The bootstrap analyses were performed with 1000 replicates, simple taxon addition and TBR branch swapping. The Consistency Index (CI), the Retention Index (RI) and the Homoplasy Index (HI) were calculated from the consensus tree (Electr. Suppl.: Table S1).

PartitionFinder (Lanfear & al., 2012) was used to find the best model of evolution and the best partitioning scheme under the Bayesian information criteria (BIC; Schwarz, 1978), for purpose of the Bayesian Inference (BI) analyses. All loci were defined as unique partitions and the models tested were those implemented in BEAST and MrBayes. A greedy search algorithm was selected for running the analysis for each dataset. The BI analysis of the cpDNA sequences was conducted with MrBayes v.3.2 (Ronquist & al., 2012). For the analysis of the coded indels of the rpl32 the simplest possible model, i.e. the Jukes Cantor model, was used. We generated 10,000 trees running MrBayes for 5,000,000 generations and sampling one of every 500 generations. After ensuring that the Markov chain Monte Carlo (MCMC) reached stationarity, we discarded the first 2500 trees as burn-in.

### Divergence time estimation. –

The dating analysis was performed using the nrDNA sequences because of the low resolution obtained with the cpDNA gene tree and also with a nrDNA-cpDNA combined analysis (not shown), and the incongruences found between the two genomes. After a preliminary analysis using all the specimens sampled (Electr. Suppl.: Fig. S1), we pruned the data set to include only one specimen for each taxon to represent only the cladogenetic events that resulted in speciation or different genetic lineages. Accordingly, for *C. aequitriloba* (Viv.) A. Chev., we included three individuals representing three genetic lineages: *Cymbalaria aequitriloba* 1 represented the Corsican lineage; *C. aequitriloba* 3, the Balearic lineage; and *C. aequitriloba* 5, the Sardinian lineage. Due to the absence of fossils in the tribe, we performed the analysis with a secondary calibration point obtained from Vargas & al. (2013). This approach has been criticized but is accepted in the absence of fossils for a direct calibration (Forest, 2009). The use of the nrDNA gene tree for the divergence time estimation analyses could also cause phylogenetic artefacts derived from the concerted evolution of ribosomal genes (Soltis & al., 1998; Álvarez & Wendel, 2003). Using a fully resolved, multi-locus phylogeny would be desirable (Maddison & Knowles, 2006), but molecular dating based on nrDNA has been successfully used in cases of low resolution and high level of polymorphism of cpDNA markers and incongruence with nrDNA markers (e.g., Gao & al., 2015; Nie & al., 2015).

The dating analysis was performed in BEAST 2 (Bouckaert & al., 2014), and the divergence between *Epixiphium wilizeni* (A. Gray) Munz and the clade *Asarina procumbens* Mill.-*Cymbalaria* was modelled as a normal distribution with a mean of 20.8 Ma and a standard deviation of 4.4 Myr. A birth-death model was employed as a tree prior (Gernhard, 2008). We used an uncorrelated relaxed clock lognormal model of the lineage-specific rate variation (Drummond & al., 2006) and set a uniform distribution with ranges of 5x10^−4^–5×10^−2^ substitutions per site per Ma (s/s/Ma, Blanco-Pastor & al., 2012). These constraints include the previous estimates for hebaceous plants ITS (1.7–8.3×10^−3^ s/s/Ma; Kay & al., 2006) and ETS rates (1.3–2.4 fold higher than ITS rates; Baldwin & Markos, 1998). Two MCMCs were run for 100,000,000 generations, and the trees were sampled every 10,000 generations. The details of the model are in the.xml file (available on request from the corresponding author). We verified the convergence of runs and that stationarity was reached with the inspection of effective sample sizes in Tracer v1.6.0 (Rambaut & al., 2013). The trees were combined with LogCombiner v1.6.2 after discarding the first 25% of the trees as burn-in. We summarized the output in a maximum clade credibility (MCC) tree with TreeAnnotator v1.6.2.

To represent diversification through time, we used the R-package APE 3.3 (Paradis & al., 2004) to construct the lineage-through-time (LTT) plots. We used the dAICRC test (Rabosky, 2006a) as implemented in the R-package LASER (Rabosky, 2006b) to infer whether the diversification rate changed over time. We tested the observed value of dAICRC against a null distribution of dAICRC values obtained from 1000 random phylogenetic trees generated under the constant rate pure birth model. The MCC tree and a random sample of 1000 trees from the posterior distribution from the dating analysis were used as input files after pruning the outgroup taxa.

### Ancestral-area estimation. –

We used the dated tree after pruning the outgroup taxa, since neither *Asarina* nor *Epixiphium* shared their distribution with any *Cymbalaria* taxon and a preliminary analysis including the outgroup resulted in very ambiguous estimation of ancestral areas for the basal nodes (not shown). We considered eight areas (Fig. 2) based on previously defined biogeographic patterns (Takhtajan, 1986; Rivas-Martínez & al., 2004) and on the endemism and distribution patterns of *Cymbalaria*.

**Figure 2.**
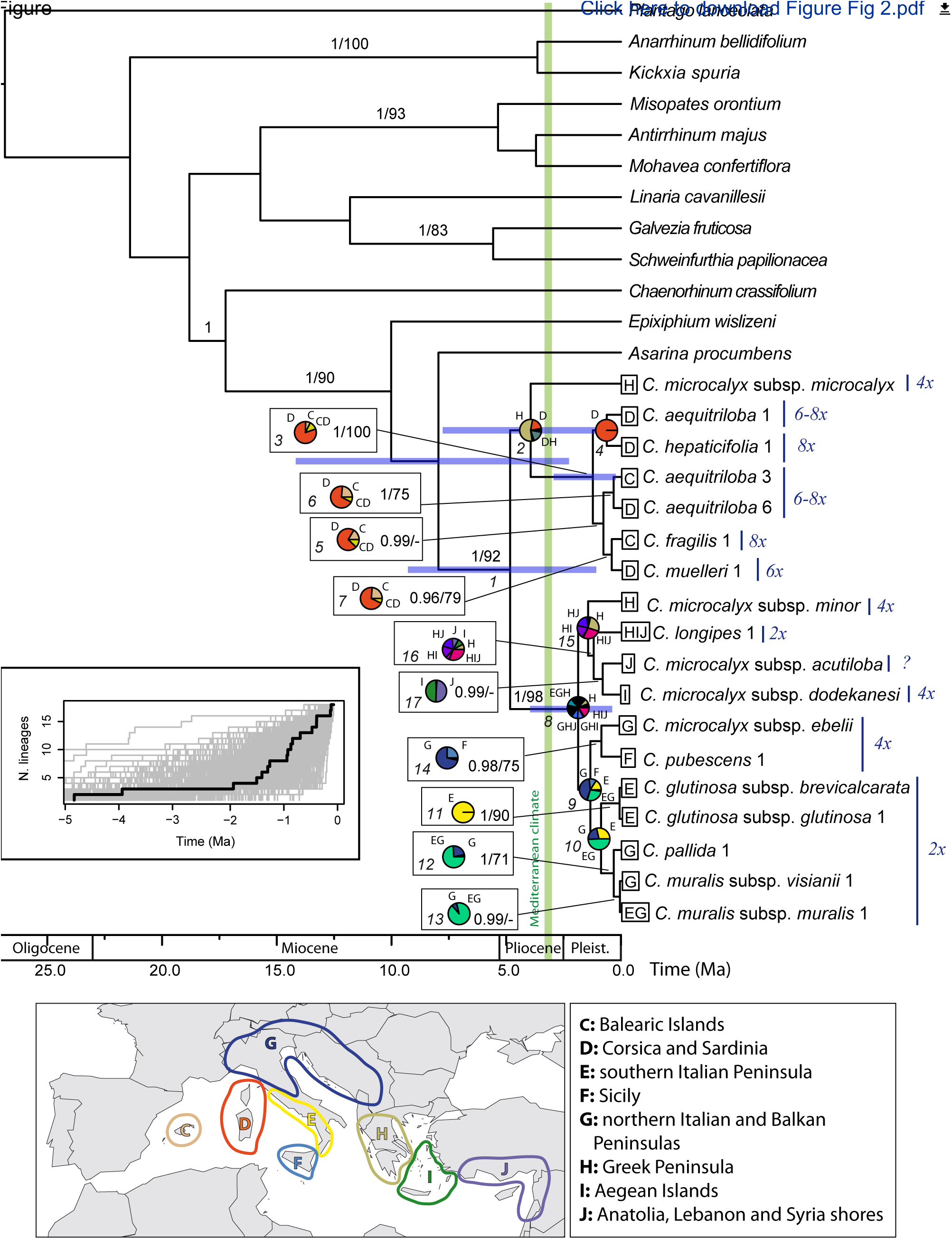
Maximum clade credibility (MCC) tree produced with a relaxed molecular clock analysis of the nrDNA for the genus *Cymbalaria* in BEAST2. Node bars represent the 95% highest posterior density intervals for the divergence time estimates of the clades that are discussed in the main text. Bayesian posterior probabilities ≥ 0.95/bootstrap support values ≥ 70% are indicated. Ploidy levels are indicated to the right of the terminal names. The numbers in italics under nodes indicate the node number. Pie charts at each node show the marginal probabilities of alternative ancestral ranges obtained from the BioGeoBEARS analysis and are shown only for the nodes discussed in the main text. Letter codes for each area inferred and distribution areas at present are indicated in the nodes and terminals, respectively. Black in pie charts represents ancestral ranges with a probability < 5%. The inset shows a lineage-through-time plot for *Cymbalaria*, based on 1000 trees randomly sampled from the posterior distribution of the dating analysis of data set 3. The thick line corresponds to the MCC tree.

We performed the biogeographic analysis with BioGeoBEARS (Matzke, 2013). This R-package implements six biogeographic models in a common likelihood framework: a likelihood version of Dispersal-Vicariance analysis (DIVALIKE; Ronquist, 1997), LAGRANGE Dispersal and Extintion Cladogensis (DEC) model (Ree & al., 2005; Ree & Smith, 2008), a likelihood version of BayArea (Landis & al., 2013), and an alternative version for each of the models that includes founder-event speciation (+J). BioGeoBEARS has two primary advantages compared with other biogeographical programs: 1) the best model is selected with likelihood ratio tests, and 2) founder-event speciation is included, a process ignored by most other methods.

The maximum number of areas for each node was set to 3, which is the maximum number of areas occupied by extant taxa (Ronquist, 1996; Hilpold & al., 2014). Each terminal in the tree was coded with the total distribution area of the taxon/lineage, except for *C. muralis*, that was only coded for its natural distribution area. We defined a dispersal probability matrix to determine the effect of geographic distance on dispersal ability. The rate of dispersal between western (Fig. 2; C, D) and eastern Mediterranean areas (H, I, J) was set to 0.5 following Hilpold & al. (2014) and was set to 1 for the other cases, to reflect the low probabilities of dispersing from eastern to western Mediterranean areas without an intermediate stage in the central Mediterranean areas (E, F, G). We ran the six models and after testing them with a likelihood ratio test and the Akaike Information Criterion (AIC), the DEC+J model was selected.

## RESULTS

### Phylogenetic analyses. –

The analyses of the nrDNA with MP and BI resulted in congruent topologies (Fig. 2, Electr. Suppl.: Table S1, Fig. S1). *Cymbalaria* was recovered as a monophyletic genus (Fig. 2, PP = 1; BS = 92%) sister to *A. procumbens*, and these two genera together subsequently sister to *E. wislizeni* (PP = 1; BS = 90%). Two main lineages were obtained within *Cymbalaria*, respectively composed of the central and eastern Mediterranean species (centre-east lineage, PP = 1; BS = 98%) and the western Mediterranean species (west lineage, PP = 1; BS = 100%). *Cymbalaria microcalyx* (Boiss.) Wettst. subsp. *microcalyx* was sister to the west lineage without statistical support (PP = 0.55).

The analyses of the cpDNA with MP and BI resulted in a congruent topology with each other (Fig. 3, Electr. Suppl.: Table S1). *Cymbalaria* was monophyletic (PP = 1; BS = 77%) and grouped with *A. procumbens* (PP = 1; BS = 76%) and these two genera with *E. wislizeni* (PP = 1; BS = 95%). The phylogenetic position of *Chaenorhinum crassifolium* (Cav.) Lange was incongruent with the nrDNA analyses, but congruent with previous cpDNA phylogenies (Ghebrehiwet & al. 2000; Vargas & al., 2013). Resolution at the species level was lower compared to the nrDNA analyses and a few incongruences were detected. In the cpDNA analysis *C. microcalyx* subsp. *ebelii* (Cufod.) Cufod. grouped with *C. glutinosa* Bigazzi & Raffaelli, *C. muralis* and *C. pallida* Wettst., (PP = 0.99) while in the nrDNA analyses it formed a clade with *C. pubescens* (J. Presl & C. Presl) Cufod (Fig. 2, PP = 0.98; BS = 75%). Slightly incongruent phylogenetic relationships were obtained in the western lineage too. For the taxa with two or more specimens sampled, only *C. glutinosa* subsp. *glutinosa*, *C. pallida* and *C. pubescens* were monophyletic in both the cp and nrDNA data sets (Fig. 3, Electr. Suppl.: Fig. S1).

**Figure 3.**
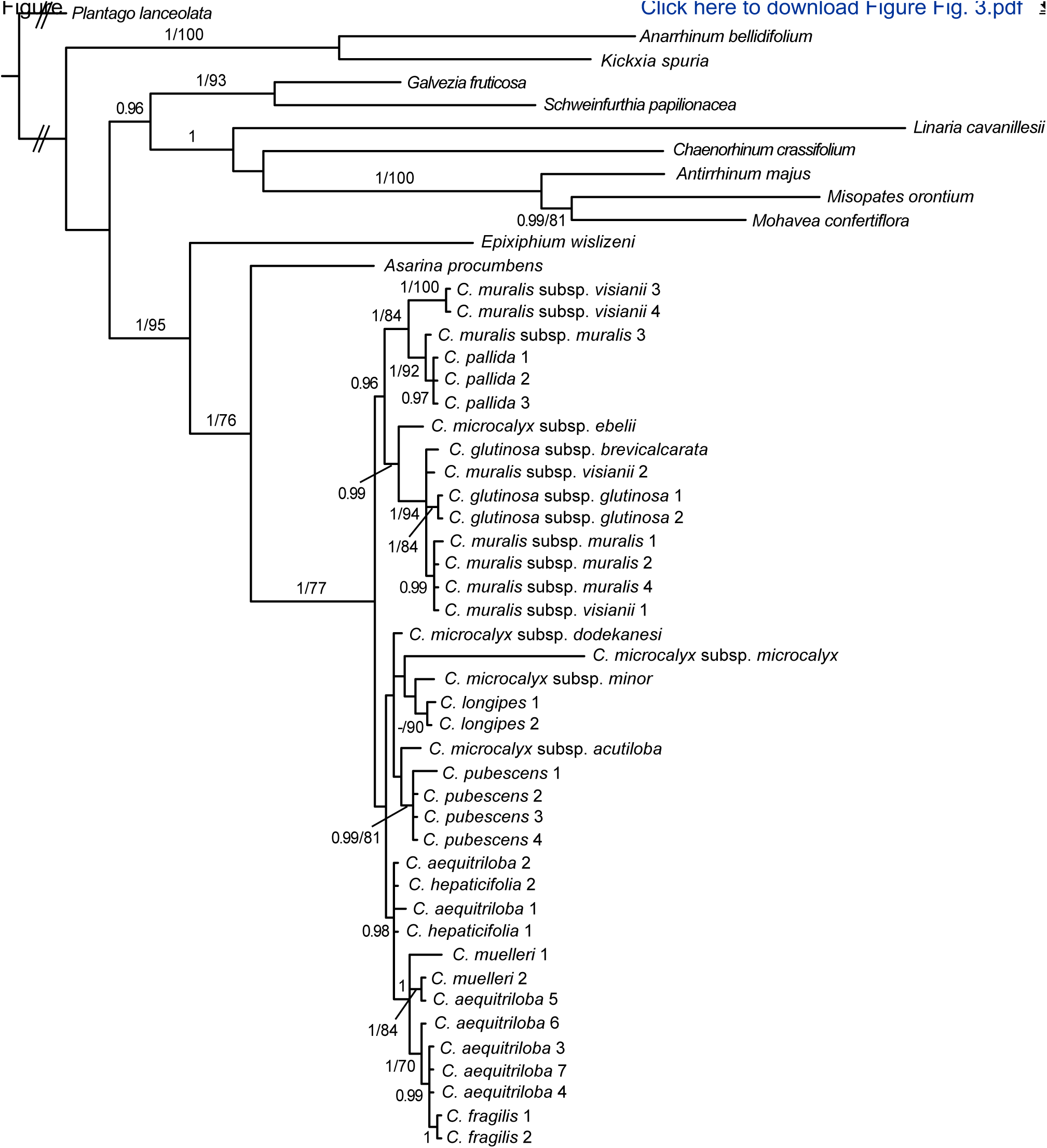
Phylogram from the Bayesian analysis of the cpDNA for the genus *Cymbalaria*. Bayesian posterior probabilities ≥ 0.95/bootstrap support values ≥ 70% are indicated. The double slashes at the base of the tree indicate that respective branches have been manually shortened.

### Divergence time estimation analyses. –

The first diversification within *Cymbalaria* occurred 4.83 Ma (node 1, 9.28–1.07 Ma 95% HPD). The first cladogenesis events in the centre-east and west lineages occurred 1.87 Ma (node 8, 3.96–0.38 Ma 95% HPD) and 1.23 Ma (node 3, 2.93–0.22 Ma 95% HPD), respectively. The LTT plot showed a notable increase of diversification towards the present. However, the dAICRC test did not reject the null hipothesis of a constant rate of diversification (p-value = 0.99). Among the two constant rate models tested, a pure birth model was selected (dAICRC = −1.31).

### Ancestral-area estimation. –

Many different areas were recovered with similar probability values with the DEC+J analysis for the ancestral area of the MRCA of *Cymbalaria* (not shown). The MRCA of the west lineage was most probably distributed in Corsica-Sardinia (Fig. 2, node 3, P = 81%), and two dispersal events to the Balearic Islands were inferred within this clade in nodes 6 and 7. Several areas with similar probability values were recovered for the ancestral area for the centre-east lineage MRCA (node 8), the eastern species clade (nodes 15 and 16) as well as for the basal nodes of the central species clade (9 and 10). The MRCA of *C. pubescens* and *C. microcalyx* subsp. *ebelii* was probably distributed in the northern Italian and Balkan peninsulas (node 14, P = 69%) and a subsequent dispersal to Sicily originated *C. pubescens. Cymbalaria pallida* and *C. muralis* subsp. *visianii* (Jáv.) D.A. Webb evolved in the northern part of the distribution of their MRCA, which occupied the Italian and northern Balkan peninsulas (nodes 12 and 13, P = 75% and 92%, respectively).

## DISCUSSION

### The origin of *Cymbalaria* and early diversification. –

Based on our results, *Cymbalaria* split from *Asarina* in the late Miocene-Pliocene (Fig. 2). Although statistical supports were low for this node in the nrDNA analyses, *Asarina* was also found to be sister to *Cymbalaria* with high statistical support in the cpDNA analysis (Fig. 3), as well as in previous studies with both cp and nrDNA (Ghebrehiwet & al., 2000; Vargas & al., 2004, 2013). Several ancestral areas with low probability values were estimated for the MRCA of *Cymbalaria*. However, a western origin can most probably be discarded, given that low ploidy levels are found in the central and eastern Mediterranean basin (Fig. 1), and western taxa are all polyploids, thus presumably of more recent origin (e.g. Garcia-Jacas & Susanna, 1992).

The low resolution observed in the basal nodes of *Cymbalaria* in the nrDNA tree and the LTT plot might reflect a rapid initial diversification (Riina & al., 2013; Vitales & al., 2014), followed by an apparent lack of diversification until ca. 2 Ma, when diversification increased rapidly (Fig. 2). However, the dAICRC test could not reject a constant rate of diversification since the origin of the genus. The pattern of increased diversification towards the present observed in the LTT plot could be explained by the “pull of the present” phenomenon (Nee & al., 1994; Kubo & Iwasa, 1995). A constant extinction rate can result in an excess of recently diverged lineages that could lead to the wrong conclusion of an increase of the diversification rate (Nee & al., 2001). This phenomenon is also the reason why detecting increases in the diversification rate is more difficult than decreases, and therefore results should be interpreted with caution (Rabosky, 2006a).

### The establishment of the Mediterranean climate and the diversification of lineages. –

The diversification of the two observed lineages occurred after the onset of the Mediterranean climate (3.2 Ma, Fig. 2), supporting the role of this climatic event as a trigger for diversification within many Mediterranean plant lineages (Fiz-Palacios & Valcárcel, 2013, and references therein). In this particular case, the Mediterranean climate likely favoured isolation of *Cymbalaria* populations in small, suitable microclimatic areas, favouring allopatric speciation events.

Ancestral area for the centre-east lineage was poorly estimated, and its descendant east and central subclades were low statistically supported, but they are congruent with the primary genetic barrier found in other groups (e.g. *Cerastium dinaricum* Beck & Szyszy?.: Kutnjak & al. 2014; *Edraianthus graminifolius* A.DC.: Surina & al., 2014) and with the floristic provinces (Takhtajan, 1986). The decline in global temperature during Pleistocene glaciations (2.6–0.01 Ma, Thompson, 2005; Médail & Diadema, 2009) would have presumably restricted the suitable habitats for *Cymbalaria* in the continent to only small patches along the coast on the northern Balkan Peninsula. This scenario could explain the origin of the barrier between the eastern and central areas on both sides of the Balkans, which points to an allopatric speciation process. For the central Mediterranean species, three supported clades were recovered. The clades *C. glutinosa* (Fig. 2, node 11) and *C. muralis-C. pallida* (Fig. 2, node 12) represented two groups of diploid taxa with partially sympatric distribution areas and unequal ecological requirements: *Cymbalaria glutinosa* occurs in warm Mediterranean areas in the southern half of Italian peninsula, whereas *C. pallida* and *C. muralis* occupy northern, wetter and cooler places in the Italian peninsula that extend to the northern Balkan peninsula in the case of *C. muralis* (Pignatti, 1982; Fig. 1, Table 1). In the same line, whereas *C. muralis* occupies humid lowlands, *C. pallida* is endemic to the highest elevations of the Apennine Range (Pignatti, 1982; Fig. 1, Table 1). In both cases, sympatric speciation was inferred. The third clade was composed by the tetraploids *C. pubescens* and *C. microcalyx* subsp. *ebelii* (Fig. 2, node 14). Their common ancestor was inferred to be present on the northern Italian and Balkan peninsulas, from which a dispersal to Sicily and further isolation originated *C. pubescens*, a route also proposed for other plant groups (e.g. *Centaurea cineraria* L. group: Hilpold & al., 2011; *Edraianthus graminifolius*: Surina & al., 2014). However, according to the cpDNA phylogeny (Fig. 3), *C. microcalyx* subsp. *ebelii* would be closely related to central Mediterranean taxa, but not to *C. pubescens*, which instead was grouped with eastern taxa without statistical support. The eastern Mediterranean clade was not supported statistically at its basal node (Fig. 2, node 15), and the ancestral area estimation was ambiguous recovering the highest probabilities for ancestors distributed across two to three eastern Mediterranean areas. The Aegean Islands were reconnected many times to the mainland during the Pleistocene climatic oscillations, which led to range expansions and subsequent allopatric speciation events when the sea level increased (Polunin, 1980). Founder-event speciation was inferred for the split between *C. microcalyx* subsp. *dodekanesi* Greuter and subsp. *acutiloba* (Boiss. & Heldr.) Greuter (Fig. 2, node 17), although the direction of the dispersal event was not clear. By contrast, the fluctuations in sea level had no apparent effect on *C. longipes* (Boiss. & Heldr.) A. Chev.; this species is widely distributed on coastal cliffs of the Aegean region with apparent adaptations to marine dispersal (Sutton, 1988), which would lead to a continuous gene flow.

A Corso-Sardinian origin for the west lineage during the Pleistocene was supported (Fig. 2, node 3). Founder-event speciation was reconstructed for *C. fragilis* (J.J. Rodr.) A. Chev. after a longdistance dispersal (LDD) event from Corsica-Sardinia (Fig. 2, node 7). At least one more LDD event was inferred for the range expansion of *C. aequitriloba* to the Balearic Islands (Fig. 2, node 6). These two areas were last connected approximately 20 Ma (Speranza & al., 2002), and therefore, a vicariant alternative to the LDD event (suggested by Verlaque & al., 1993) must be discarded. Long-distance dispersal events were previously invoked to explain the origin of some of the endemic plant species with a disjunct Balearic-Corso-Sardinian distribution (e.g. *Thymus herba-barona* Loisel.: Molins & al., 2011). Moreover, Nieto Feliner (2014) reports that LDD events have not been rare in the Mediterranean, even when no particular adaptations for seed dispersal exist. The success in colonization of new areas is often linked more to pre-adaptations of genotypes and availability of suitable habitats than to distance (Alsos & al., 2007). Polyploidy may have been a key trait in the colonization processes because it potentially provided an increased ability to tolerate a wide range of ecological conditions (Ramsey, 2011). Additionally, because of low interspecific competition in rocky habitats and low diversity on islands, more niches would be available, with the likely consequence of higher success in colonization (Thompson, 2005).

### Speciation. –

The three primary types of speciation likely occurred throughout the evolution of *Cymbalaria*. Based on our results combined with published chromosome counts, the hypotheses of allopatric speciation, sympatric speciation and polyploid speciation are supported.

Allopatric speciation is inferred when sister taxa occupy different areas isolated by physical barriers. The two main types of allopatric speciation are vicariance and founder-event speciation, which results from founder events. In historical biogeography, vicariance has long been recognized as a key process in diversification (Ronquist, 1997), and implies that a widely distributed ancestor gives rise to two or more separate species within its original distribution area when the appearance of a physical barrier provokes their reproductive isolation. However, in *Cymbalaria*, vicariance was not inferred for any statistically supported node. By contrast, founder-event speciation is a process largely ignored in historical biogeographical models but is currently recognized as an essential process (Gillespie & al., 2012; Matzke, 2013). It involves a rapid divergence of a small, peripheral population of a species, and is inferred when the area of one of the descendants is not part of the ancestor’s distribution area. Indeed, the selection of the DEC+J model indicated that founder-event speciation (parameter J) was important for the model to fit our data. Our results supported founder-event speciation in three cases: the origin of *C. pubescens* (Fig. 2, node 14), the split between *C. microcalyx* subsp. *acutiloba* and *C. microcalyx* subsp. *dodekanesii* (Fig. 2, node 17), and the origin of *C. fragilis* (Fig. 2, node 7). This last case shows the typical structure of a founder-effect speciation event, where the newly and rapidly generated species (*C. fragilis*) is embedded in a more widely distributed and genetically variable, paraphyletic species (Futuyma, 2005), in this case *C. aequitriloba* (Fig. 2). For *C. fragilis*, LDD was inferred (see above), but in the other two cases, the low sea levels during the Pleistocene glaciation periods may have favoured a stepping stone dispersal to new areas (e.g., Campanulaceae: Cellinese & al., 2009; *Centaurea cineraria* group: Hilpold & al., 2011).

DEC+J model inferred sympatric speciation in six statistically supported clades (Fig. 2, nodes 3, 4, 5, 11, 12 and 13). However, geographical and ecological isolation are not mutually exclusive, and their effects are difficult to disentangle from one another (Papadopulos & al., 2014). Most of the inferred cases of sympatric speciation in our results could be interpreted as artefacts of the resolution used when defining the areas. For example, the split between the Corsican lineage (*C.aequitriloba* 1 and *C. hepaticifolia* 1) and the rest of the taxa within the west lineage (Fig. 2, node 3) might have been a case of geographical isolation of this island from Sardinia. Moreover, geographical isolation can also occur at a local scale, particularly for plants that grow in rock crevices such as *Cymbalaria*. These habitats can be scarce and isolated from one another, favouring small-scale allopatric speciation processes (Thompson, 2005). However, in groups of taxa where gene flow is possible due to long distance dispersal, the recognition of putative geographic barriers is a difficult task. An additional impediment is that distribution areas can change over time, and current sympatric species could have originated allopatrically and later expanded their areas to become sympatric. Apart from these limitations, to infer sympatric ecological speciation, it would be desirable to demonstrate that adaptation to the different ecological niches exists and it is actually the cause of reproductive isolation, assuming that ecological niches have not changed significantly from the speciation moment until the present (Carine & Schaefer, 2009). Very local scale environmental measures would be required to properly describe ecological niche in the case of *Cymbalaria*, since the habitats where they occur (Table 1) have usually very different microclimatic conditions than the general climatic available data, which make methods such as species distribution modelling fail (Guisan & Thuiller, 2005; Austin, 2007). In our group of study, sympatric ecological speciation could explain the differentiation of *C. muralis* and *C. pallida*, as inferred by the DEC+J model (Fig. 2, node 12). These two species occur in the same region (northern Italy), often within a few hundreds of metres of one another (P. Carnicero & M. Galbany-Casals, personal observations) and occupy different niches (Table 1). However, their distributions at local scale are almost allopatric, given that *C. muralis* mostly occupies the lowlands while *C. pallida* grows in higher elevations. Thus, allopatric speciation could not completely be ruled out.

The important role of polyploid speciation in the diversification of *Cymbalaria* was already suggested by others (Verlaque & al., 1993; Thompson, 2005). Biogeographic analyses never integrate polyploid speciation; however, we consider that genome duplication event occurred when a supported clade composed by specimens of the same ploidy level is found (as assumed by e.g. Wood et al., 2009, Marcussen et al., 2015), and therefore that the clade originated from a polyploid speciation event. Polyploid species placed in different supported clades would have had been originated by independent polyploidization events. Accordingly, two polyploid speciation events are hypothesized: one for the origin of the west lineage (Fig. 2, node 3) and the other for the origin of the -*C. microcalyx* subsp. *ebelii-C. pubescens* clade (Fig. 2, node 14). The monophyly of the west lineage species apparently refutes the hypothesis of independent polyploid origins for *C. hepaticifolia* Wettst. and *C. aequitriloba* from the diploids *C. pallida* and *C. muralis* of the Italian Peninsula, respectively, as suggested by Verlaque & al. (1993). However, the polyploid clades could be the result of interlocus concerted evolution of the nrDNA (ITS and ETS), which may hide the genetic information of one of the parental lineages in the case of allopolyploids (Wendel & al., 1995). This hypothesis has to be specially considered for the clade *C. microcalyx* subsp. *ebelii* – *C. pubescens*, given that these two species appear in separate clades in the cpDNA analysis (Fig. 3). Additional studies are required to confirm the common origin of the species in each of the two polyploidy clades and to distinguish between the auto-and allopolyploidization events and the parental taxa involved. The support to LDD events found here for the western clade (see subsection: The establishment of the Mediterranean climate and the diversification of lineages) is consistent with the observed pattern of higher probability of LDD events in polyploid groups (Linder & Barker, 2014). This pattern may be associated to the high genetic variability of polyploids but also to their difficulty in succeeding in areas in which the parental species occur (Thompson, 2005; Ramsey, 2011).

### Conclusions. –

Many characteristics of the genus *Cymbalaria* are of value to increase our understanding of the processes that shaped the current biodiversity of the Mediterranean. The evolution of *Cymbalaria* was marked by climatic events, particularly by the onset of the Mediterranean climate and the Pleistocene climatic oscillations. Both geographical and ecological barriers played important roles in the speciation of the genus, but the identical barrier might have had disparate consequences for closely related taxa, as shown by two examples in both the eastern and western Mediterranean. In both cases, one species is widely distributed across sea (*C. aequitriloba* in the west and *C. longipes* in the east), whereas the same geographic isolation acted as barrier and originated *C. fragilis* and the differentiation of *C. microcalyx* subspecies, respectively in the western and eastern Mediterranean Basin. Polyploidy, by increasing the ability to colonize new areas and promoting rapid speciation, is proposed as a key process in the diversification of *Cymbalaria*. The supported monophyly of the genus found for both the cp and nrDNA analyses support its current taxonomy status. However, our study revealed some conflicts between current taxonomy and phylogeny at the species and subspecies level, being *C. microcalyx* the most shocking case, with some of its subspecies showing very distant phylogenetic positions from each other (Figs. 2, 3). When more than one specimen per taxon was used, only *C. glutinosa* subsp. *glutinosa*, *C. pallida* and *C. pubescens* were monophyletic for both the cp and nrDNA (Fig. 3, Electr. Suppl.: Fig. S1). Detailed studies combining molecular and morphological data are needed to definitely unravel the taxonomy of *Cymbalaria*.

## ACKNOWLEDGEMENTS

We thank all herbaria that provided material and colleagues who provided assistance during fieldwork or their own plant material. We also thank M. Fernández-Mazuecos for providing the data on the secondary calibration point for the dating analyses and C. Roquet for technical assistance on the ancestral-area estimation. This research was funded in part by the Spanish government (CGL2010-18631/BOS and *Flora iberica* project, CGL2011-28613-C03-01) and the Catalan government (2009-SGR 439 and 2014-SGR 514). Pau Carnicero benefited from the support of a PIF Ph.D. student fellowship from the Universitat Autònoma de Barcelona.

## Appendix 1.

Sampled specimens with information on the individual numeric codes used in text and figures, locality, herbarium voucher and accession numbers of the regions analysed. A dash (–) indicates sequences that were not obtained in the present study or specimens without individual numeric code.

Taxon, individual number, locality, voucher, GenBank acc. no. ITS, 3’ETS, *ndhF*, *rpl32-trnL*

***Cymbalaria*** Hill: ***C. aequitriloba*** (Viv.) A. Chev., 1, France, Corsica, La Castagniccia, *A. Curcó* (BCN 86695), KP735225, KP851084, KP851014, KP851100; ***C. aequitriloba***, 2, France, Corsica, La Castagniccia, *A. Hilpold s. n*. (BOZ 8888), KP735224, KP851085, KP851011, KP851097; ***C. aequitriloba***, 3, Spain, Balearic Islands, Mallorca, Puig Major, *X. Rotllan* (no voucher), KP735219, KP851088, KP851007, KP851093; ***C. aequitriloba***, 4, Spain, Balearic Islands, Mallorca, Formentor, *L. Sáez 7366 & X. Rotllan* (BC), KP735240, KP851086, KP851009, KP851095; ***C. aequitriloba***, 5, Italy, Sardinia, Nuoro, Badde Salighes, *C. Aedo 9213* (MA 708824), KP735220, KP851087, KP851026, KP851111; ***C. aequitriloba***, 6, Italy, Sardinia, Cuglieri, Mte. Ferru, *C. Navarro 4683 & al*. (MA 708259), KP735222, –, KP851006, KP851092; ***C. aequitriloba***, ***7***, Spain, Balearic Islands, Cabrera, *L. Sáez 6196 & L. Guárdia Valle* (BC), KP735241, KP851082, KP851008, KP851094; ***C. fragilis*** (J. J. Rodr.) A. Chev., 1, Spain, Balearic Islands, Menorca, Barranc d’Algendar, *P. Carnicero 346 & M. Galbany-Casals* (BC), KP735211, KP851081, KP851004, KP851090; ***C. fragilis***, 2, Spain, Balearic Islands, Menorca, Barranc d’Algendar, *P. Carnicero 346 & M. Galbany-Casals* (BC), *–*, –, KP851005, KP851091; ***C. glutinosa*** Bigazzi & Raffaelli subsp. ***glutinosa***, 1, Italy, Spigno Saturnia, *P. Carnicero 734 & M. Galbany-Casals* (BC), KP735216, KP851068, KP851029, KP851114; ***C. glutinosa*** subsp. ***glutinosa***, 2, Italy, Spigno Saturnia, *P. Carnicero 734 & M. Galbany-Casals* (BC), KP735217, KP851069, KP851030, KP851115; ***C. glutinosa*** subsp. ***brevicalcarata*** Bigazzi & Raffaelli, Italy, Ravello, *P. Carnicero 748 & M. Galbany-Casals* (BC), KP735218, KP851070, KP851020, KP851105; ***C. hepaticifolia*** Wettst., 1, France, Corsica, Lac du Nino, *A. Hilpold s. n*. (BOZ 8842), KP735223, KP851079, KP851022, KP851107; ***C. hepaticifolia***, 2, France, Corsica, Castagniccia, *P. Carnicero 444 & M. Galbany-Casals* (BC), KP735215, KP851078, KP851013, KP851099; ***C. longipes*** (Boiss. & Heldr.) A. Cheval., 1, Greece, Dodecanese Islands, Karpathos, *N. Böhling 8228* (B 100138948), KP735232, KP851064, KP851038, KP851123; ***C. longipes***, 2, Greece, Samos, *E. Gathorne-Hardy 657* (E 629368), *–*, –, KP851039, KP851124; ***C. microcalyx*** (Boiss.) Wettst. subsp. ***microcalyx***, Greece, Peloponnese, Lakonia, *W. Greuter & H. Merxmüller s. n*. (B 100460657), KP735238, KP851063, KP851041, –; ***C. microcalyx*** subsp. ***acutiloba*** (Boiss. & Heldr.) Greuter, Turkey, Antalia, Alanya, *P. H. Davis 25847 & O. Polunin* (E 629362), KP735212, KP851059, KP851042, KP851126; ***C. microcalyx*** subsp. ***dodekanesi*** Greuter, Greece, Rhodes, Archangelos, *P.H. Davis 40310* (E 629364), KP735208, KP851058, KP851043, KP851127; ***C. microcalyx*** subsp. ***ebelii*** (Cufod.) Cufod., Montenegro, Skadar Lake, *E. Mayer 11192 & M. Mayer* (B 100460658), KP735236, KP851061, KP851036, KP851121; ***C. microcalyx*** subsp. ***minor*** (Cufod.) Greuter, Greece, Kefallinia, Aenos, *J. Damboldt s. n*. (B 100460655), KP735237, KP851060, KP851037, KP851122; ***C. muelleri*** (Moris.) A. Chev., 1, Italy, Sardinia, Seui, Genni d’Acca, *P. Carnicero 406 & M. Galbany-Casals* (BC), KP735210, KP851080, KP851012, KP851098; ***C. muelleri***, 2, Italy, Sardinia, Ulassai, *P. Carnicero 389 & M. Galbany-Casals* (BC), KP735209, KP866214, KP851010, KP851096; ***C. muralis*** G. Gaertn., B. Mey. & Scherb. subsp. ***muralis***, 1, Spain, Catalonia, Sant Cugat (naturalized), *P. Carnicero* (no voucher), KP735230, KP851077, KP851015, KP851089; ***C. muralis*** subsp. ***muralis***, 2, Spain, Catalonia, Caldes de Montbui (naturalized), *P. Carnicero 135* (BC), KP735231, KP851076, KP851017, KP851102; ***C. muralis*** subsp. ***muralis***, 3, Poland, Slask Dolny (naturalized), *Z. Pulawska s. n*. (FI),*–*,–, KP851018, KP851103; ***C. muralis*** subsp. ***muralis***, 4, Italy, Toscana, Albegna, *F. Selvi s. n*. (FI), *–*, –, KP851019, KP851104; ***C. muralis*** subsp. ***visianii*** (Jáv.) D. A. Webb, 1, Italy, Lazio, Palombara, *P. Carnicero 703 & M. Galbany-Casals* (BC), KP735226, KP851075, KP851027, KP851112; ***C. muralis*** subsp. ***visianii***, 2, Italy, Lazio, Palombara, *P. Carnicero 703 & M. Galbany-Casals* (BC), *–*, –, KP851028, KP851113; ***C. muralis*** subsp. ***visianii***, 3, Italy, Lazio Rocca di Papa, *P. Carnicero 710 & M. Galbany-Casals* (BC), KP735226, KP851074, KP851031, KP851116; ***C. muralis*** subsp. ***visianii***, 4, Italy, Lazio Rocca di Papa, *P. Carnicero 710 & M. Galbany-Casals* (BC), *–*, –, KP851032, KP851117; ***C. pallida*** Wettst., 1, Italy, Abruzzo, Valle d’Orfenta, *P. Carnicero 780 & M. Galbany-Casals* (BC), KP735234, KP851072, KP851033, KP851118; ***C. pallida***, 2, Italy, Abruzzo, Valle d’Orfenta, *P. Carnicero 780 & M. Galbany-Casals* (BC), KP735235, KP851071, KP851034, KP851119; ***C. pallida***, 3, Italy, Abruzzo, l’Aquila, *J. Aldasoro 3276* (MA 698766), KP735233, KP851073, KP851035, KP851120; ***C. pubescens*** (J. Presl & C. Presl) Cufod., 1, Italy, Sicily, Palermo, La pizzuta, *C. Aedo 5733 & al*. (MA 646152), KP735229, KP851066, KP851021, KP851106; ***C. pubescens***, 2, Italy, Sicily, Trapani, Erice, *J. Güemes 3085 & al*. (SALA 106642), KP735214, KP851065, KP851024, KP851108; ***C. pubescens***, 3, Italy, Sicily, Trapani, Mt. Acci, *C. Aedo 5614 & al*. (MA 646631), KP735228, KP851067, KP851025, KP851110; ***C. pubescens***, 4, Italy, Sicily, Trapani, Mt. Acci, *J. Güemes 3052 & al*. (SALA 106608), KP735213, –, KP851023, KP851108; Other Antirrhineae: *Anarrhinum bellidifolium* (L.) Willd., Spain, Catalonia, l’Espluga de Francolí, *M. Galbany-Casals 2303* (BC), KP735199, –, KP851052, KP851136; ***Antirrhinum majus*** L., Spain, Catalonia, Alella, *M. Galbany-Casals 2302* (BC), KP735205, –, KP851048, KP851132; ***Asarina procumbens*** Mill., Spain, Catalonia, Montseny massif, *P. Carnicero 253 & L. Sáez* (BC), KP735207, KP851057, KP851045, KP851129; ***Chaenorhinum crassifolium*** (Cav.) Lange, Spain, Valencian Country, Serra d’Aitana, *P. Carnicero 207 & al*. (BC), KP735203,–, KP851051, KP851135; ***Epixiphium wislizeni*** (A. Gray) Munz, USA, New Mexico, Animas Valley, *G.R. Ballmer s. n*. (RSA 712541), KP735206, KP851056, KP851046, KP851130; ***Galvezia fruticosa*** J. F. Gmel., Perú, Lima, Yauyos, *M. Weigend 7209 & al*. (B 100095831), KP735197, –, KP851044, KP851128; ***Kickxia spuria*** (L.) Dumort. subsp. ***integrifolia*** (Brot.) R. Fern., Spain, Catalonia, Gallecs, *J.M. Blanco s. n*. (BC), KP735200, –, KP851053, KP851137; ***Linaria cavanillesii*** Chav., Spain, Valencian Country, Dènia, *P. Carnicero 197 & al*. (BC), KP735198, –, KP851050, KP851134; ***Misopates orontium*** (L.) Rafin., Spain, Valencian Country, Fenestrat, *P. Carnicero 210 & al*. (BC), KP735201, –, KP851049, KP851133; ***Mohavea confertiflora*** A. Heller, USA, California, Colorado desert, *T.R. Stoughton 800* (RSA 778206), KP735202, –, KP851054, KP851138; ***Plantago lanceolata*** L., Spain, Catalonia, Cerdanyola, *P. Carnicero 523* (BC), KP735196, –, KP851055, KP851139; ***Schweinfurthia papilionacea*** Boiss., Oman, Nizwa, *A. G. Miller 6657* (E 614757), KP735204, –, KP851047, KP851131.

